# Brain-based Lifespan Trajectories of Social Cognition: From Resting-state fMRI Perspective

**DOI:** 10.1101/2022.07.22.501083

**Authors:** Wei Fan, Zhi-Xiong Yan

**Affiliations:** Key Laboratory of Brain and Behavior Science, Nanning Normal University, Nanning 530023, China; Department of Psychology, Hunan Normal University, Changsha 410000, China

**Author notes:** Corresponding Author: Zhixiong Yan, Ph.D.

**Keywords:** Lifespan trajectories, Social Cognition, Lifespan Trajectories, Intrinsic functional connectivity

## Abstract

Brain lifespan trajectories of intrinsic functional connectivity (iFC) in social cognition (SC) provide comprehensive insight and effective reference for normal development and clinical intervention. Unfortunately, for a lack of lifespan samples, the neural mechanism underlying SC in a lifespan developmental perspective is largely unknown, though relative behavior evidence has been investigated well. We, based on common functional networks, aim to map out voxel-wise iFC features between the SC networks and the entire cerebral cortex and further chart its corresponding trajectory patterns across the lifespan. Three networks, namely moral cognition, theory of mind (ToM), and empathy, are chosen as representatives of core SC networks. NKI-RS Enhanced (N = 316, ages 8-83 years old) dataset is selected as a lifespan resting-state fMRI dataset to delineate iFC characteristics and construct developmental trajectories. The result shows that the SC networks represent dissociable and network-specific connectivity profiles. Empathy network, as the most age-related susceptible network, shows a linear up trend with dorsal attention network, a linear decrease with ventral attention network, and a quadric-concave with somatomotor and dorsal attention networks. In addition, the sex effect is also discovered in empathy network, which exhibits linear and quadric gender differences with frontoparietal and vision networks respectively. ToM network follows a pronounced quadric-convex (inverted U-shape) trajectory with ventral attention network. No linear and quadric trajectory is found in moral cognition network. These findings indicate that SC networks display a wide range of iFC both with the fundamental networks (e.g., somatomotor, vision) and advanced networks (e.g., attention and control) in specific developmental trajectories, which will provide a better understanding of SC neural mechanism and life-span developmental trajectories also give valuable references for SC abnormal detection and effective therapy.

## Introduction

Social cognition (SC) supplies fundamental supporting for better understanding and interaction with each other in human society (Burnett, Sebastian, Kadosh, & Blakemore, 2011; F. Happé, Cook, & Bird, 2017). Some unitized structures may subserve SC stimuli, such as medial prefrontal cortex (MPFC), posterior cingulate cortex (PCC), temporoparietal junction (TPJ), and superior temporal sulcus (STS) (Adolphs, 2001), which are mainly located in default mode network (DMN), a typical intrinsic network playing a critical role in social cognition (Raichle, 2015, for review). Previous studies investigated the neural mechanism of the structure of SC development (Mills, Lalonde, Clasen, Giedd, & Blakemore, 2014) or SC atypical development from infant to adolescence (Happe & Frith, 2014). Other evidence also focused on lifespan development with a limited age range or lack of specific SC concentration (Coupe, Catheline, Lanuza, & Manjon, 2017; Craik & Bialystok, 2006; Wang, Su, Shen, & Hu, 2012; Yang et al., 2014). To our best knowledge, SC functional connectivity development across the lifespan is still rare to challenges in debating networks definition and the entire lifespan dataset (Adolphs, 2001; Frith & Frith, 2008), which implies a promising approach from a lifespan developmental perspective to explore its neural mechanism (Zuo et al., 2016). Thanks to advances in the meta-analysis of SC networks and public-shared lifespan neuroimaging datasets (e.g., Bzdok et al., 2012; Nooner et al., 2012), it is becoming possible to construct iFC lifespan trajectories of SC networks. The present work aims to focus on three core SC networks (Empathy, Theory of Mind (ToM), and Moral cognition) (Bzdok et al., 2012) to map out its iFC patterns in common cortical networks (Yeo et al., 2011) and construct corresponding lifespan developmental trajectories through NKI-RS lifespan fMRI dataset (Nooner et al., 2012).

Empathy was thought to be a proxy for sharing interpersonal experiences and mental states (Decety, 2010, 2011). Plenty of studies using behavioral paradigms explored its age-associated features with limited age phases. Specifically, infants as young as 14 to 18 months began to develop an intention to help others attain goals (Warneken & Tomasello, 2009). This competence continued to advance unremittingly and formulated an intrinsic empathy motivation when age reaches from childhood to adolescence (Casey, Tottenham, Liston, & Durston, 2005; Fabes, Fultz, Eisenberg, May-Plumlee, & Christopher, 1989; Paus, 2005). However, from adults on, the view on empathy development was disparate. Some studies emphasized that older individuals were more efficient and had more positive empathy ratings than younger adults (Beadle, Sheehan, Dahlben, & Gutchess, 2015; F. G. Happé, Winner, & Brownell, 1998). The others reported that aging might bring a deleterious effect on empathy-related social cognitive performance (Charlton, Barrick, Markus, & Morris, 2009). An age-related decline was also reported in a functional MRI study when subjects perceived the other agent who experience pain (Chen, Chen, Decety, & Cheng, 2014). Overall, the evidence above demonstrated that empathy developed beginning in infancy period and continuing through childhood, young adulthood, and even in the elderly (Duval, Piolino, Bejanin, Eustache, & Desgranges, 2011, for a review).

The development of ToM, termed as inferring what others thought, intention, or behavior, is regarded as the fundamental underpinning for childhood SC development (Flavell, 1999; Premack & Woodruff, 1978). Substantial studies supposed that the course of ToM development proceeded along infants and extended to adults. Specifically, a 12 mouths child could share attention (Leung & Rheingold, 1981), while children at age two began to distinguish between pretense or desires (Henry M Wellman & Woolley, 1990) and performed false belief tasks successfully around 5 years old (Perner & Wimmer, 1985; Zaitchik, 1990), which might increasingly be protracted to early adolescence and adulthood (Rieffe, Terwogt, & Cowan, 2005; Rosenberg-Kima & Sadeh, 2010; Williams et al., 2009) with more efficient and dedicate cognitive development (Brunet, Sarfati, Hardy-Bayle, & Decety, 2000; Maylor, Moulson, Muncer, & Taylor, 2002). With aging, The unimpaired ToM performance in a specific domain might be through compensating mechanism to activate or strengthen more brain structure or functional connectivity for elders than younger adults (Amodio & Frith, 2006), while impaired ToM in social aging is associated with a significant decline in task performance such as executive functioning (for review in Moran, 2013). Taken together, the capacity of ToM strengthened from children to elders, but this evidence is based more on behavior observation than neural mechanism investigation, which is hotly debated and needs to warrant in future research.

Moral cognition, a more comprehensive component in SC, refers to judging others’ right-wrong intentions or behaviors. Its development attracted numerous investigations with discrete findings across the lifespan. The most influential moral development theory may attribute to Kohlberg’s cognitive stages development, which proposed that moral cognition developed from lower to higher levels based on the progress of fundamental cognitive development (Kohlberg, 1971). Recently, numerous developmental studies suggested that moral cognition might follow an accumulating developing trend. Moral-like intentions (e.g., preferentially interacting with pro-social agents) would be robust for 6-months infants (Hamlin, Wynn, & Bloom, 2007). 14–18-month-old infants know how to comfort a distressed person and could exhibit unrewarded helping behavior to others(Warneken & Tomasello, 2009). Identifying the notions of truth from lie and shift judgmental standards from outcome to intention would facilitate the development of pre-mature moral judgment in children around 8-year-old (Cushman, Sheketoff, Wharton, & Carey, 2013; Lăzărescu, 2012). With experience accumulation, early adolescence (aged 11-14 years) could independently disengage moral cognition from actual moral behavior, nearly reaching moral maturity (Caravita, Sijtsema, Rambaran, & Gini, 2014). In addition, a task-fMRI study has reported that age-related activation decreases in the amygdala and increases in the vmPFC when participants viewed morally salient scenarios (Decety, Michalska, & Kinzler, 2012). A relatively stable pattern with intertwined tendencies has also been reported from preschool to 31-32 years old (Lapsley & Carlo, 2014). Those results showed that the developmental trend of moral cognition might be complex and need to wait for methodological innovation to parse its chronologic variability.

Above all, much behavioral evidence (albeit inconsistent) about SC development was accumulated with the limited investigation of its neuro mechanism. It is necessary to apply a lifespan neuroimaging dataset to parsing SC neural mechanisms and better understand iFC of SC development.

## Methods

### Participants

The research was reviewed and approved by the ethics committee of Nanning Normal university. The resting-state fMRI lifespan data used in this study are based on publicly available datasets, NKI-Rockland (NKI-RS) enhanced, which consist of 316 healthy participants (mean age 44.38, 112 males) over the age range 8-83 years(Nooner et al., 2012)(Pic. X for details). All other participants with psychiatric conditions are excluded. The dataset usage approved by NKI-RS review institution. All participants provided informed written consent before participating in NKI-RS experiments.

### Image preprocessing

The pipeline of Connectome Computation System (CCS: http://github.com/zuoxinian/CCS)(Ting Xu, 2015) was used for preprocessing images, mapping individual, group-level connectomics, trajectory constructions and visualization with exceling performance on the cortical mantle.

Structural MRI data preprocessing primarily includes (1) denoising, skull stripping and structural segmentation through Volbrain, an automatic and online brain volumetry system (Manjon & Coupe, 2016), (2) cortical surface reconstructing by FreeSurfer (http://www.freesurfer.net), (3) boundary-based registration (BBR) for alignment (Greve & Fischl, 2009). The rfMRI image preprocessing mainly incorporates (1) denoising and removing motion artifacts through ICA-AROMA (Pruim et al., 2015); (2) removed of the first 8 volumes to reach stable scanning; (3) masked by the functional brain mask generated based on both structural brain and 4D individual time series; (4) normalized to have a comparable 4D global mean intensity of 10,000 across individuals; (5) regressed out of mean signals from white matter and cerebrospinal fluid masks as well as Friston-24 head motion parameters, intracranial volume, and errBBR; (6) temporally smoothed with a band-pass filtering; (7) detrended by removing both linear and quadratic trends in time domain; and (8) projected onto the fsaverage surface grid and down-sampled to the fsaverge5 surface grid.

### Functional connectivity and trajectories construction

We used the dual regression approach to build individual-level DR components (Beckmann, Mackay, Filippini, & Smith, 2009; Filippini et al., 2009; Zuo et al., 2010).

This method is based on the following GLM dual regression equations:

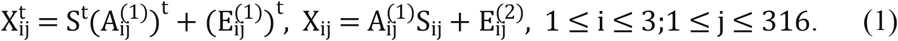

In Eq. (1), X_ij_ represents the fMRI data from the i-th social network of the j-th participant. The first part of Eq. (1) uses unthresholded group-level components S as the spatial predictors of the individual fMRI volumes and results in the regression matrix A_ij_ containing the relevant individual regression weights in ij the time domain (i.e., timeseries). These timeseries were then used as the temporal predictors for the individual fMRI timeseries in the second regression equation. The resulting regression matrix Sij contains regression weights for each of the components in the spatial domain, which serve as our measure of network-level functional connectivity (i.e., the individual-level DR components). These individual-level DR components were subsequently used to evaluate the lifespan developmental trajectory of the group-level components after RFT correction.

To improve the reliability of trajectories, we apply a semi-parametric regression model (Generalized Additive Models for Location, Scale and Shape: GAMLSS) to construct trajectories(Rigby, Stasinopoulos, Heller, & Voudouris, 2014) as a way of overcoming some of the limitations associated with the popular Generalized Linear Models (GLM) and Generalized Additive Models (GAM).

## Results

Vertex-wise iFC and its lifespan changes between SC networks and common functional networks were analyzed. All SC networks presented significant age-related variance except moral cognitive network. Among these SC networks, the empathy network was the most complicated with both linear and inverted U-shape lifespan trajectories (Fig. 2, 3). Specifically, a linear ascending iFC trend was found between the empathy network and dorsal attention network, and a linear decline between the empathy network and ventral attention network (Fig. 2). Moreover, A quadratic-convex (inverted U-shape) iFC trend was discovered with somatomotor and dorsal attention network (Fig. 3). Moreover, sex effect was also observed between empathy network and frontoparietal control network in which male had more strengthening iFC than female across the lifespan (Figure 4). An interaction between gender and an age-related quadratic trend was further found between the empathy network and visions network (Figure 4), which show male fit a quadratic-concave (U-shape) pattern while female fit a quadric-convex (inverted U-shape) trend. Besides, the age-related changes in ToM network only exhibited an inverted U-shape trajectory with left and right ventral attention networks across the lifespan (Figure 5).

**Fig. 1.**
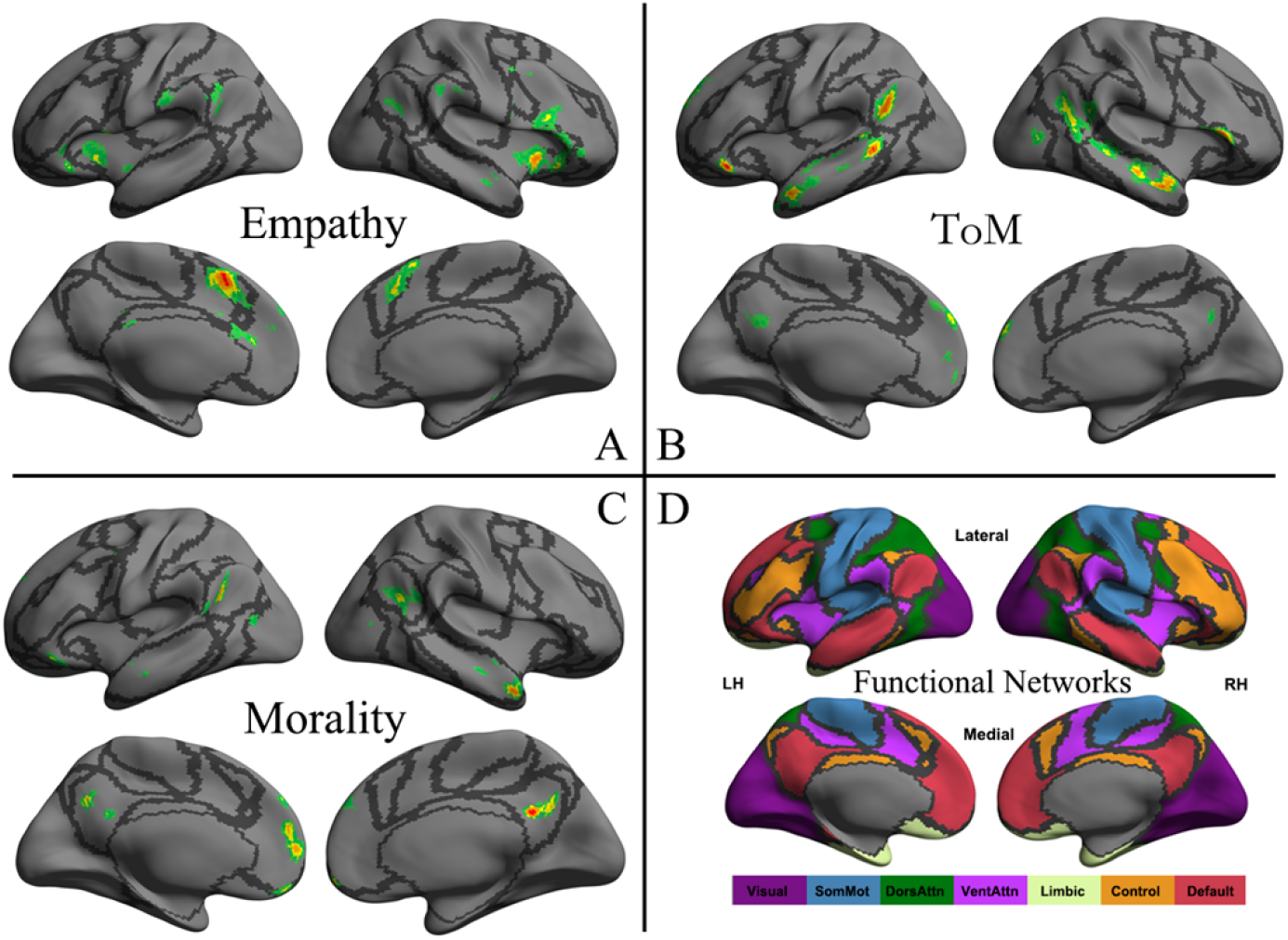
the network distribution of three social cognitive networks (A, B and C) and seven intrinsic functional connectivity networks (D). Social cognitive networks are namely empathy network (A), ToM network (B) and moral cognitive network (C). Seven intrinsic functional networks established from Yeo et al. (2011) functional parcellation based upon 1,000 healthy participants including visual (Visual), somatomotor (SomMot), dorsal attention (DorsAttn), ventral attention (VentAttn), limbic (Limbic), frontoparietal control (Control), and default mode (Default) networks. Each panel plots in left (LH) and right (RH) hemisphere and lateral (Lateral) and medial (Medial) view respectively. The cortical grid model “inflated_pre” of the fsaverage in FreeSurfer was employed. Gray curves represent the boundaries of the seven networks.

**Fig. 2.**
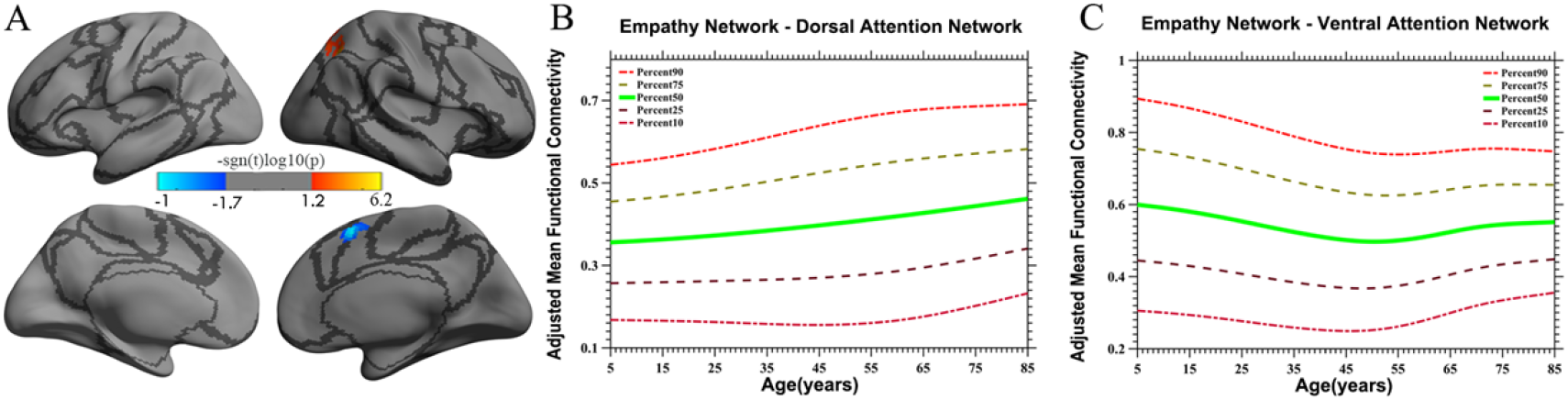
The linear lifespan iFC pattern of empathy network located in dorsal attention network (red in A) and Ventral attention network (blue in A). The surface distribution (A) was estimated by general linear model (GLM) analysis and RFT correction in freesurfer and CCS. The lifespan trajectories are constructed by a semi-parametric regression models: Generalized Addictive Models for Location, Scale and Shape (GAMLSS). The color bar in panel A means linear increasing trend (red, plot in B) and linear decreasing (blue, plot in C). The 5 lines in panel B and C represents 90, 75, 50, 25 and 10 percentile line respectively.

**Fig. 3.**
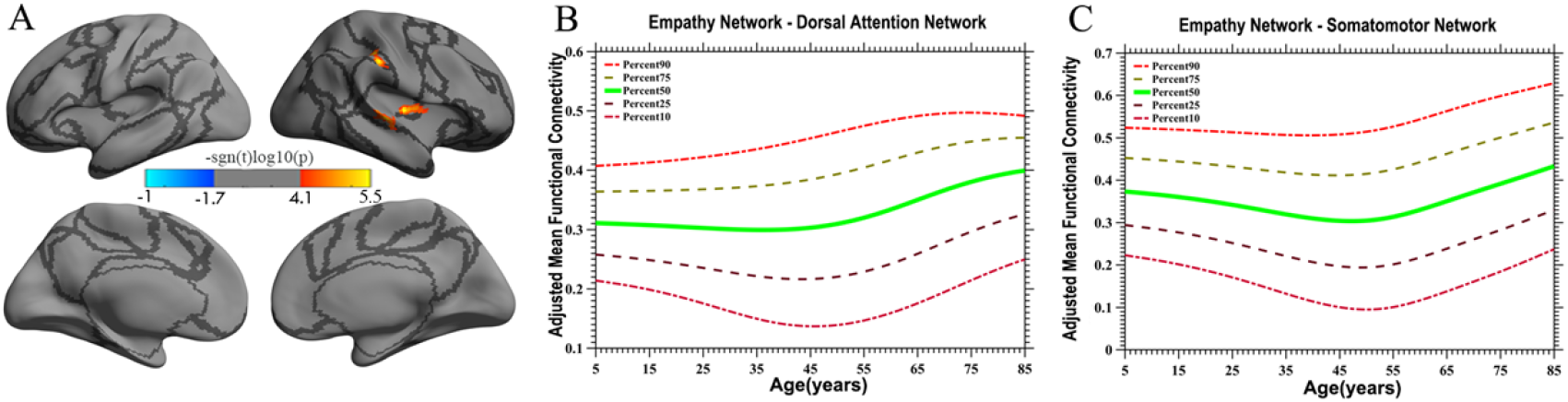
The quadric lifespan iFC pattern of empathy network located in dorsal attention network and Somatomotor network. The surface distribution (A) was estimated by general linear model (GLM) analysis and RFT correction in freesurfer and CCS. The lifespan trajectories (B, C) are constructed by a semi-parametric regression models: Generalized Addictive Models for Location, Scale and Shape (GAMLSS). Red color (A) located in dorsal attention network and somatomotor network means quadric increasing trend (U-shape) which plot in panel B and C respectively. The 5 lines in panel B and C represents 90, 75, 50, 25 and 10 percentile line respectively.

**Fig. 4.**
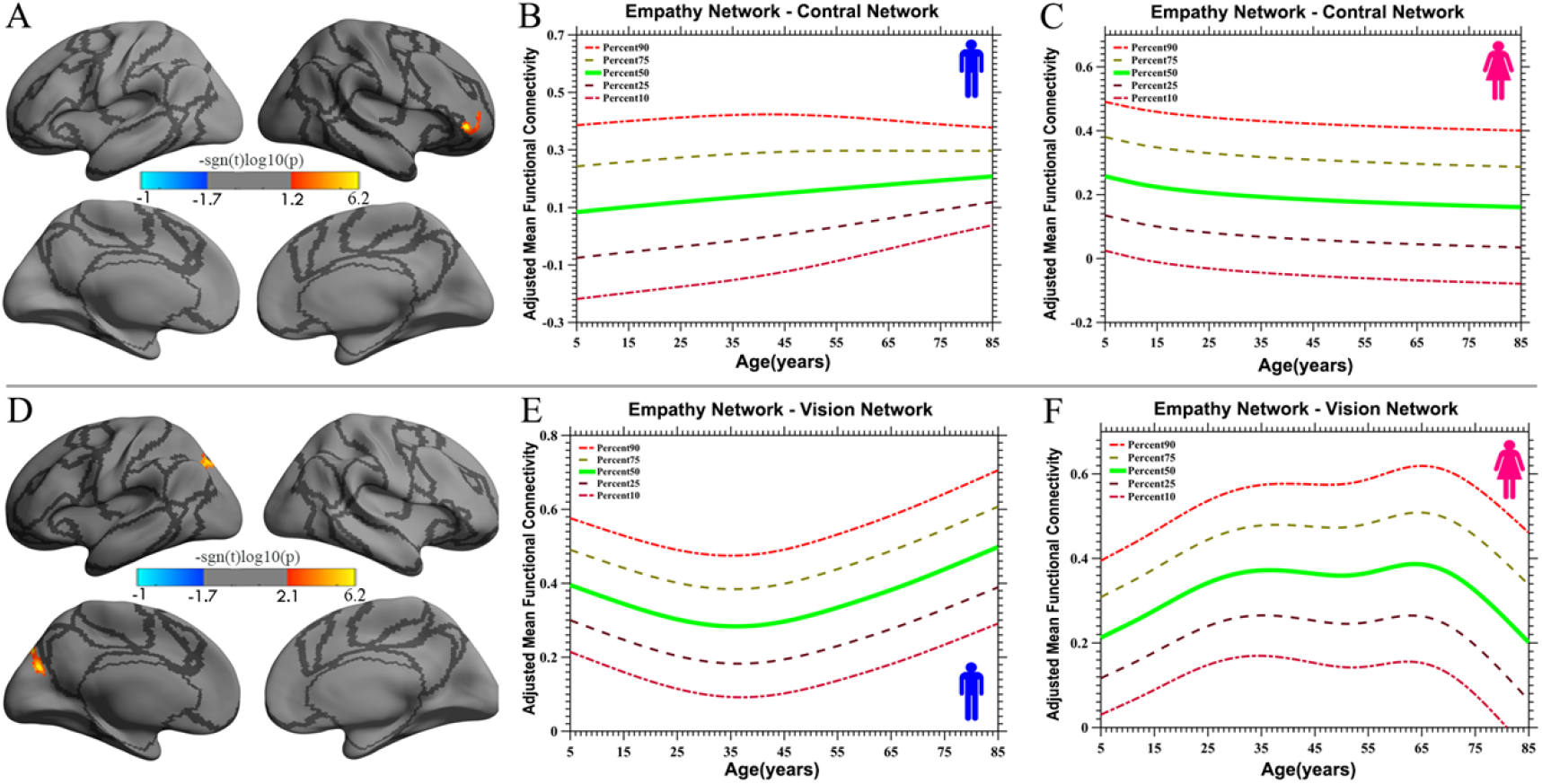
The lifespan iFC trajectories of gender effects of empathy network which located in control network (linear, A) and vision network (quadric, D). The surface distribution (A, D) was estimated by general linear model (GLM) analysis and RFT correction in freesurfer and CCS. The lifespan trajectories (B, C, E, F) are constructed by a semi-parametric regression models: Generalized Addictive Models for Location, Scale and Shape (GAMLSS). Panel B, E and C, F indicate the trend of male and female trajectories respectively. The 5 lines in panel B, C, E and F represent 90, 75, 50, 25 and 10 percentile line respectively.

**Fig. 5.**
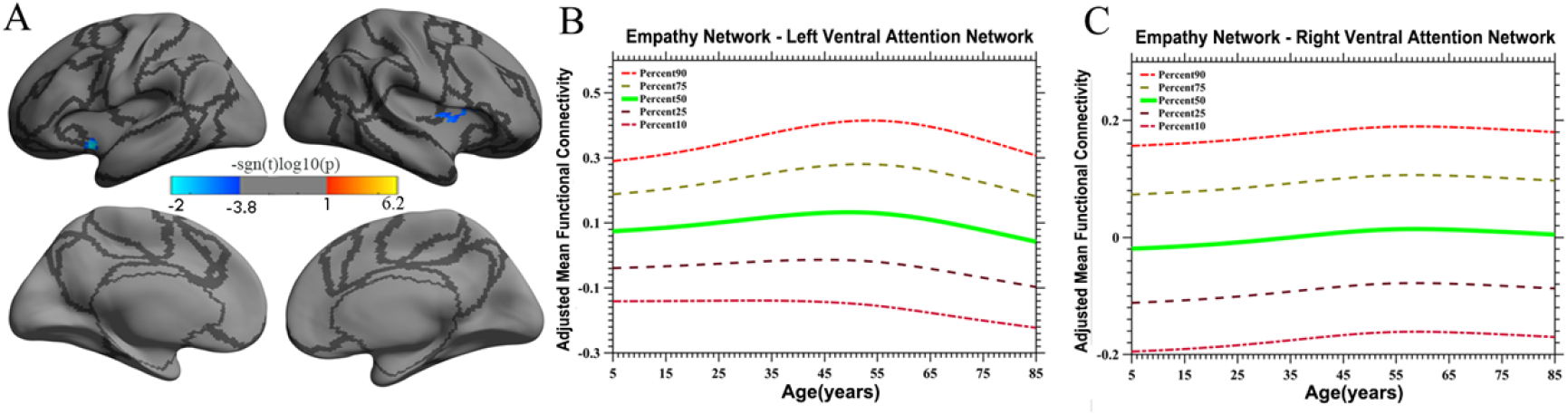
The lifespan quadric iFC trajectories of ToM network which located in left and right ventral attention network. The surface distribution (A) was estimated by general linear model (GLM) analysis and RFT correction in freesurfer and CCS. The lifespan trajectories (B, C) are constructed by a semi-parametric regression models: Generalized Addictive Models for Location, Scale and Shape (GAMLSS). Panel B and C plot the trajectory located in left and right ventral attention network respectively. The 5 lines in panel B and C represent 90, 75, 50, 25 and 10 percentile line respectively.

**Table 1.** Sample characteristics and MRI acquisition protocol

|  | NKI – RS _Pilot | NKI – RS _Enhanced | CCNP |
| --- | --- | --- | --- |
| **Demography** |  |  |  |
| Mean Age (SD) | 36.84(21.20) years | 44.38(19.72) years | 11.91(3.32) years |
| Range | 7-85 years | 8-83 years | 6.53-17.91 years |
| N | 126 | 316 | 185 |
| Male/Female | 68/58 | 112/204 | 86/99 |
| **Scanner** |  |  |  |
| Manufacturer | Siemens | Siemens | Siemens |
| Tesla | 3.0 | 3.0 | 3.0 |
| **MP-RAGE** |  |  |  |
| TR | 2,500 ms | 1,900 ms | 2,600 ms |
| TE | 3.50 ms | 2.52 ms | 3.20 ms |
| Inversion time | 1,200 ms | 900 ms | 1,200 ms |
| Flip angle | 8° | 9° | 8° |
| Field of view | 256 mm | 250 mm | 256 mm |
| Slices | 192 | 176 | 176 |
| Voxel size | 1.0 × 1.0 × 1.0 mm | 1.0 × 1.0 × 1.0 mm | 1.0 × 1.0 × 1.0 mm |
| **EPI** |  |  |  |
| TR | 2,000 ms | 645 ms | 2,500 ms |
| TE | 30 ms | 30 ms | 30 ms |
| Flip angle | 75° | 60° | 80° |
| Field of view | 224 mm | 222 mm | 216 mm |
| Slices | 34 | 40 | 38 |
| Voxel size | $3.5 \times 3.5 \times 3.5$ mm | $3.0 \times 3.0 \times 3.0$ mm | $3.0 \times 3.0 \times 3.3$ mm |
| Number of slices | 147 | 900 | 186 |

## Discussion

We discovered the three SC networks had extensive iFC with the cerebral cortex networks (vision, somatomotor, ventral attention, dorsal attention, and control network) and showed specific trajectory features across the lifespan (linear and quadric trends). The iFC of the empathy network showed richer age-related variations (linear increase and decrease, inverted U-shape) with dorsal attention network and somatomotor network than ToM and moral network. The iFC of ToM network was presented in a quadratic-convex trajectory with ventral attention network. A gender effect was also found in the empathy network. From the above, we postulated that SC might manifest distinct developmental profiles and functional connectivity properties across the lifespan.

Large-scale functional networks have been becoming an attractive tool to explore the underlying neuronal mechanism. Several rs-fMRI studies demonstrated that the lifespan development of functional networks showed a relatively stable way across the entire lifespan, though the microstructure and function of the brain might likely be variable (Fair et al., 2009; Fransson, Aden, Blennow, & Lagercrantz, 2011). This evidence provided a possibility to probe iFC characteristics of SC network with other functional networks and to construct developmental trajectories across the entire lifespan. Its neurobiological application of these observations would be extensive for normal developmental monitoring and clinical intervention and evaluation.

### iFC trajectory of empathy network across lifespan

The iFC of the empathy network, as mentioned above, played the most dramatic and distinct development profiles including both linear and nonlinear trajectories. A linearly increasing trend was found with dorsal attention network (Fig. 2). Previous studies demonstrated that dorsal attention network was involved in top-down regulation (Fox, Corbetta, Snyder, Vincent, & Raichle, 2006; Koziol, Barker, Joyce, & Hrin, 2014). The increasing iFC with dorsal attention network may mean that human empathy development has a continuous strengthening top-down monitoring with accumulating experience, which was inconsistent with the study of Greimel et al. (2010) who found an age-related increased activity in the fusiform gyrus and inferior frontal gyrus that mainly located in control network. The reason for this discrepancy may be the limited age range (aged 8-27 years) that is not sufficient to match the lifespan developmental profile as in the present study. A study of empathy-related behavior supported that the older adult group who did not show impaired state-cognitive empathy but scored higher on subtests of state-affective empathy did more emotional empathy behavior relative to the younger group, indicative of an enhancing top-down mediation in older ages (Ze, Thoma, & Suchan, 2014). This result needs more direct evidences in future research.

We also found a linearly decreasing iFC trend with ventral attention network. Ventral attention network was mainly implicated in stimulus-driven attentional control (Vossel, Geng, & Fink, 2014) and made key contributions to down-up processing (for review in Etkin, Egner, & Kalisch, 2011). The pronounced decreasing iFC pattern in our results would demonstrate that empathy-related down-up processing capability followed a downstream profile, which indicated that external stimuli might lose their preponderance with growth. Chen et al. (2014) confirmed our results and showed an age-related decline FC between the anterior insula and anterior mid-cingulate cortex, partly overlaying the area discovered in the present study. Taken together, an increased top-down and decreased down-up processing pattern may reflect distinct age-related variations and specific cognitive strategies when conveying empathy to others.

Another prominent finding was that a quadric-concave (U-shape) age-related change of empathy network iFC with somatomotor network and the inferior portion of dorsal attention network (Fig. 3A), which meant this quadric-concave trend might mainly associate with sensory-motor function and top-down processing. Accumulating evidence pointed to somatosensory regions (e.g., postcentral cortex) as likely one of the candidates for encoding social perception (Meyer, Kaplan, Essex, Damasio, & Damasio, 2011; Morrison, Tipper, Fenton-Adams, & Bach, 2013). These results showed that somatosensory regions might encode social-related stimuli, which was consistent with the view of the mirroring system for socially derived information (Keysers & Gazzola, 2010). Our study extended previous literature and indicated more distinct and variable developmental characteristics with more efficient iFC with somatosensory and ventral part of dorsal attention networks in adults (40-50 years old) than children and elder, though the meaning of functional connectivity is still an open question (Power, Fair, Schlaggar, & Petersen, 2010),

Intriguingly, we further observed a gender-specific difference in iFC between the empathy network and functional networks across the lifespan. Specifically, a linear developing trend was detected between the empathy network and control network in which males presented a linear increasing trend while females presented the opposite. A quadric curving was also found with vision network where males presented a U-shaped pattern and females presented an inverted U-shaped trajectory. Numerous studies investigated gender differences in empathy development and most of them indicated that females were more empathic than males (Christov-Moore et al., 2014). Those studies mainly focused on behavior and self-report results. Other neuroimaging studies detected no sex-related differential changes in hemodynamic responses (Michalska, Kinzler, & Decety, 2013). The limited participants and specific experiment paradigms might contribute to those results. In the present study, we based on meta-analysis results and extended participants to a wide lifespan range (from 7 to 85). A linear gender-difference iFC of empathy network with control network in the present study showed that males have increasing iFC relative to female across the lifespan. Moreover, an interaction between sex and quadric curve trend find in empathy network iFC with vision network (or occipital-parietal junction), indicative male with a quadric-concave and female with a quadric-convex iFC pattern. This result is partly inconsistent with the study of Tomasi and Volkow (2012) who indicated that females had lower functional connectivity density in vision and other sensory networks than males as aging, but the interaction between age and gender was not significant. One of the reasons for this discrepancy may cause in default mode network (DMN) focused in their study while SC networks specialized in the present study. This difference also proved that SC networks were peculiar with specific gender differences in which female adults had stronger iFC relative to child and old period with vision network across lifespan development while males were on the contrary. Together, these pronounced gender differences were not only reflected in primary sensory networks but also in advanced networks like control network.

### iFC trajectory of ToM network across lifespan

The present findings showed that the iFC of ToM network presented in an inverted U-shaped lifespan trajectory with ventral attention network. A meta-analysis study of ToM development showed a linear increasing trend of ToM ability among children of 3-10 years old (Henry M. Wellman, Cross, & Watson, 2001), which is similar to our iFC developing pattern if narrowed the age span in 3-10 years. Other behavior studies also found a similar developmental trend. Children aged 6 and 7 years passed the “belief about belief” test, and children aged 9-11 years developed complex social skills, such as recognizing a social “faux-pas” or wrong behavior. Having more chances to frequently experience self-consciousness, using social comparison as a method of self-evaluation, adolescents showed a better understanding of others’ mental states as compared to pre-adolescents (Dumontheil, Apperly, & Blakemore, 2010; Vetter, Leipold, Kliegel, Phillips, & Altgassen, 2013). Aging is associated with impairments in social understanding(Moran, 2013). Neuroimaging studies about ToM development is lacking in addition to rare recent works. A iFC of ToM network development for children and adolescents showed that the middle and inferior temporal cortex and anterior cingulate cortex (key node of ToM network) were linked to the anterior insula of ventral attention network, which may be responsible for social cognition (like ToM) (Cauda et al., 2011). Age-related brain activity in the ToM network (e.g., MPFC) shifted from the ventral to the dorsal part which may associate with the maturation of the prefrontal cortex and the associated development of cognitive functions (Moriguchi, Ohnishi, Mori, Matsuda, & Komaki, 2007; Sebastian et al., 2012). Our results extended previous findings to the entire lifespan and outlined its trajectory in an inverted U-shape. For lack of direct evidence, the explanation of this result needs to be cautious.

### iFC trajectory of moral cognition network across lifespan

Among three key SC components, moral cognition is more complex and covers more neural territory than other SC (like ToM and empathy) (Greene, 2015; Moran, 2013). Moral cognitive network distributes extensively such as vmPFC/dmPFC, precuneus, temporo-parietal junction (TPJ), PCC, temporal pole, and middle temporal gyrus (Marazziti, Baroni, Landi, Ceresoli, & Dell’osso, 2013). These structures are mainly situated at the interface of ToM and empathy network, providing both rational and emotional processing (Bzdok et al., 2012). In our results, those mixed and overlapped neural structures may neutralize specific developmental trends and result in less robust age-related iFC changes. These results were supported by a longitudinal study which showed that moral cognitive related regions (such as some of the prefrontal, temporal, and parietal cortex) were relatively stable over the lifespan (Crone & Elzinga, 2015). More sensitive fitting method and participants with a wider lifespan range will be employed in our future works.

### Applications of lifespan developmental trajectories of social cognition

Normal lifespan trajectories of SC may help to better understand the underlying process of normal brain development and to further construct SC normative models. It will provide a reference for choosing educational methods for children or teenagers and providing suitable training exercise for the elderly. Furthermore, it may provide valid clinical biomarkers to guide the early identification of social cognitive disorders and evaluate suitable intervention methods. The variable age range of SC trajectories is informative to hint at a time window of effective intervention. Based on previous studies, Adolescents are characterized as a pivotal chance to intervene for their vulnerability to psychiatric conditions and atypical brain network (Kadosh, Linden, & Lau, 2013; Somerville et al., 2013). In our finding, the mature age of SC is around 35-45 years old, which is beyond the beginning of puberty until adulthood and may be more adaptive for the intervention of SC deficits than other age periods. Taken together, charting lifespan neurodevelopment of SC could contribute to a more systematic view of SC and improve our understanding of the nature of SC, which will be helpful for healthy development, early SC abnormal identification, monitoring, and treatment. The trajectories in our work will provide a start framework for SC development and clinical intervention.

### Limitations

Our study has potential caveats needed to emphasize. The structure of SC is a comprehensive domain that encompasses many vaguely subcomponents (F. Happé et al., 2017). Here, we select three core subcomponents in elucidating the social cognition lifespan trajectories, while other elements (e.g., imitation, social exclusion) are beyond the scope of the present work. In addition, the iFC between SC networks and common functional networks is based on cerebral cortex. The functional of subcortical structures like amygdala will be emphasized in our future studies. Besides, the cross-sectional samples need to be considered which may mask within-individual change by variability inter-individuals. Age-related social cognitive changes in a cross-sectional sample may reflect cohort rather than age effects, which may play a negative role in correctly constructing lifespan developmental trajectory. Our lab is collecting longitudinal cohort fMRI data to compensate for this limitation.

## Conclusion

The current study for the first time provides a heterogeneous lifespan developmental trajectory of SC networks based on iFC at vertex-wise level. Specific age-related changing profiles are found in social cognition lifespan development. Empathy network presents a relatively more complicated profile. A both decreasingly and increasingly linear iFC trend was found with dorsal attention network and ventral attention network respectively, with a U-shaped trajectory in somatomator and dorsal attention network. Moreover, increasingly linear and U-shaped gender-specific features were found in control and vision network respectively. Besides, the lifespan profile of ToM network is demonstrated in an inverted U-shape trajectory. These crucial age-related changes extend previous findings to the entire lifespan and may provide a starting point for investigating the extraordinary versatility of the developing social brain and give support to early identification and efficient intervention of social cognitive deficits.

## Acknowledgements

The funding agents had no further role in study design, data collection and analysis, decision to publish, or preparation of the manuscript. All authors declared no conflict of interest.

## Author Contributions

Conceived and designed the experiments: ZXY. Analyzed the data and wrote the paper: WF. Edit the manuscript: ZXY.

## Notes

### Competing Interest Statement

The authors have declared no competing interest.

